# Pharmacophore design and ensemble-docking as approach to screening a large-scale database for identification of potential inhibitors of APH(3')-IIIa of *Enterococcus faecalis*

**DOI:** 10.1101/2024.12.23.630124

**Authors:** Luis Alfredo Velez, Sandra Denisse Arteaga

## Abstract

Fighting nosocomial infections represents one of the main difficulties in medical treatment units around the world. Among the bacteria with the greatest potential to cause these complications is *Enterococcus faecalis*, mainly due to the expression of aminoglycoside-modifying enzymes. In the approach adopted in this work, pharmacophore modeling followed by ensemble-docking were carried out to select potential inhibitors against the nucleotide binding site of EfAPH(3')-IIIa. The most promising ligands are all nucleotide analogues (ZINC000828691112, ZINC000029546074 and ZINC000062624753) and showed good toxicological and pharmacokinetic profiles and the ability to interact with key amino acids such as MET26, SER27 and TYR42. Taken together, these preliminary results show that nucleotide analogues may represent a promising structure for interacting with the EfAPH(3')-IIIa nucleotide-binding pocket. Further analysis will be conducted.

## 1. Introduction

Aminoglycosides are secondary metabolites first isolated from soil bacteria ^1,2^.Since their isolation, first streptomycin and dihydrostreptomycin, have been the antibiotic of choice for combating a wide range of bacterial infections ^2,3^. This is due to the natural selection of these compounds acting efficiently to compete with compounds from other bacteria and also fungi ^3,4^. This class of antibiotic acts mainly by binding to the A site of 16S ribosomal RNA ^4,5^, thus acting to combat Gram-positive and Gram-negative bacteria. And the fight against infectious diseases is one of the priorities described by the World Health Organization and the health policies of many countries, especially developing countries ^6–9^.

Among the Gram-positive species largely resistant to aminoglycosides are enterococci ^10–12^. This group of bacteria are common residents of the gastrointestinal tract of humans and also of most mammals ^12–14^. When their host experiences an imbalance in the microbiome, they are also capable of causing a variety of infections, including serious infections, most frequently among hospitalized patients ^15^. One of the most widely studied representatives of the genus is *Enterococcus faecalis*, which accounts for approximately 90% of the cases of infection inferred by the enterococcus genus ^10,11^. This group is widely involved in hospital infections in transplants, catheterization and general intensive care unit infections ^13,14^.

The most widespread mechanism of bacterial resistance to aminoglycoside administration among bacterial species is the expression of aminoglycoside-modifying enzymes (AMEs) ^16,17^. This group of enzymes is divided into three main groups: 1) N-acetyltransferases (AACs) are enzymes that modify the amino groups of aminoglycosides using acetyl coenzyme A; 2) O-nucleotidyltransferases (ANTs) are the enzymes responsible for using ATP to modify the hydroxyl groups of aminoglycosides and lastly 3) O-phosphotransferases (APHs) which modify a larger group of nucleotides to covalently alter a wide range of aminoglycosides ^16–18^.

Given the limited capacity to develop new drugs in the short term, an advantageous option is to increase the search for compounds against specific targets of clinical importance ^19–21^. This large structural universe of compounds makes it possible to explore natural products, which have been used in the pharmaceutical industry for decades ^22–24^. These compounds represent the most promising sources as possible candidates for therapeutic applications, mainly due to the wide diversity of their scaffolds, mainly due to the broad diversity of their scaffolds ^24–26^. In the present approach, a pharmacophore model was built to filter the entire ZINC20 database followed by ensemble-docking method to select potential APH(3')-IIIa inhibitors capable of interacting competitively with ATP.

## 2. Methodology

### 2.1 Pharmacophore-based virtual screening using macromolecule-ligand complexes

The 3D x-ray diffraction structures for the study were obtained from RCSB Protein Database (PDB ID: 2BKK co-crystallized with ADP ^27^), (PDB ID: 3Q2J co-crystallized with CKI-7 ^28^) and (PDB ID: 3TM0 co-crystallized with ANP ^29^).

Subsequently, these three co-crystallized structures were aligned with respect to the NBS of 3TM0 using in-house python scripts as previously described ^30^. The targeted alignment resulted in the selection of common interactions between proteins and the different ligands as a resource for the pharmacophoric model using the Pharmit web tool ^31^.

Considering the x, y and z coordinates, the pharmacophore model constructed showed the following features: two aromatic rings (50.6, 49.4 and 12. 5; 51.4, 50.7 and 11.1); two hydrophobic features (50.6, 49.4 and 12. 5; 51.4, 50.7 and 11.1); two hydrogen acceptors (50.7, 50.1 and 10.1; 51.0, 49.6 and 13.6) and one hydrogen donor (49.0, 46.2 and 9.4). The virtual screening to identify hits was conducted using the ZINC database ^32^, containing 13,127,550 ligands with 122,276,899 conformers, obtained in August 2024.

The hits resulting from the database search were ranked considering mRMSD < 2.0. These hits were then treated for search of duplicate compounds, resulting in 4,620 unique ligands, additionally hydrogen atoms were assigned to the structures at pH 7.4, the 3D coordinates were generated using default settings, followed by minimization steps of these structures with the MMFF94 force field ^33^ using the steepest descent geometry optimization, followed by conjugate gradient algorithm with default parameters using Avogadro software ^34^ and transformed into mol2, all using Open Babel-3.1.1 software ^35^.

### 2.2 Ensemble docking-based virtual screening

The ensemble docking-based virtual screening strategy, to rescoring the results of pharmacophore of the 4,620 compounds, was conducted using the Protein-Ligand ANT System-1.2 software (PLANTS-1.2) ^36^. All crystals were protonated, repairing missing residues were performed and then the structures were converted into mol2 format, all using the Dock Prep module of UCSF Chimera-1.16 ^37^. At the same time, each structure was processed by the SPORES 1.3 tool using default parameters. All the analysis were conducted using the co-crystal coordinates at the nucleotide-binding pocket of 2BKK, 3Q2J, and 3TM0, with a radius of 15 Å.

The coordinates were 10.3, 27.7, and -10.4 on the x, y, and z axes to 2BKK; 27.9, 9.6, and 70.7 on the x, y, and z axes to 3Q2J; and 51.3, 42.5 and 15.5 on the x, y, and z axes to 3TM0. To ensure effective clustering in docking processing, a cut-off value was set based on RMSD values of 2.0 Å and default settings were used for all other parameters. The ensemble docking-based virtual screening results were using the consensus strategy based on the exponential method to define the compounds with the best performance. The method was implemented using customized python scripts assisted by the Pandas and Numpy libraries. The five best scores were evaluated by its pharmacokinetic preditions and subsequently molecular dynamics analysis.

### 2.3 Toxicological and pharmacokinetic predictions

The profile of hepatotoxicity, carcinogenic, mutagenicity, and cytotoxicity was analyzed by using the ProTox-II web server ^38^. Additionally, the pharmacokinetic predictions of gastrointestinal (GI), blood-brain barrier (BBB), Glycoprotein-P (P-gp) and cytochromes (CYT) were conducted using SwissADME web server ^39^.

## 3. Results and Discussion

### 3.1 First round of virtual screening assisted by pharmacophore

Due to the low determination of active compounds capable of inhibiting the activity of the EfAPH(3')-IIIa ^40^, pharmacophore modeling based on macromolecule-ligand ^41^ crystallographic structures was initially adopted. The molecular interactions considered to generate the pharmacophore were extracted from the common interactions of the aligned structure of the co-crystals CKI-7, ANP and ADP in the nucleotide binding pocket of APH3'IIIa. It was observed that the active site has TYR42, SER91 and ALA93 as its main hydrogen bonds donor components, Fig 1A. In addition, it was also observed that the surface of the enzyme contains numerous GLU and ASP residues (ten residues), Fig 1B, giving the active site an acidic character that should be explored for a potential inhibitor interaction. Therefore, a model with two aromatic, two hydrophobic, two H-acceptor and additionally one H-donor features were contemplated, Fig. 1C. This pharmacophore design also aimed to select hits with a nucleotide core, since these compounds are established as effective against various infectious diseases like emerging viruses ^42^ and have structural similarity to the physiological ligand, ATP.

**Fig 1.**
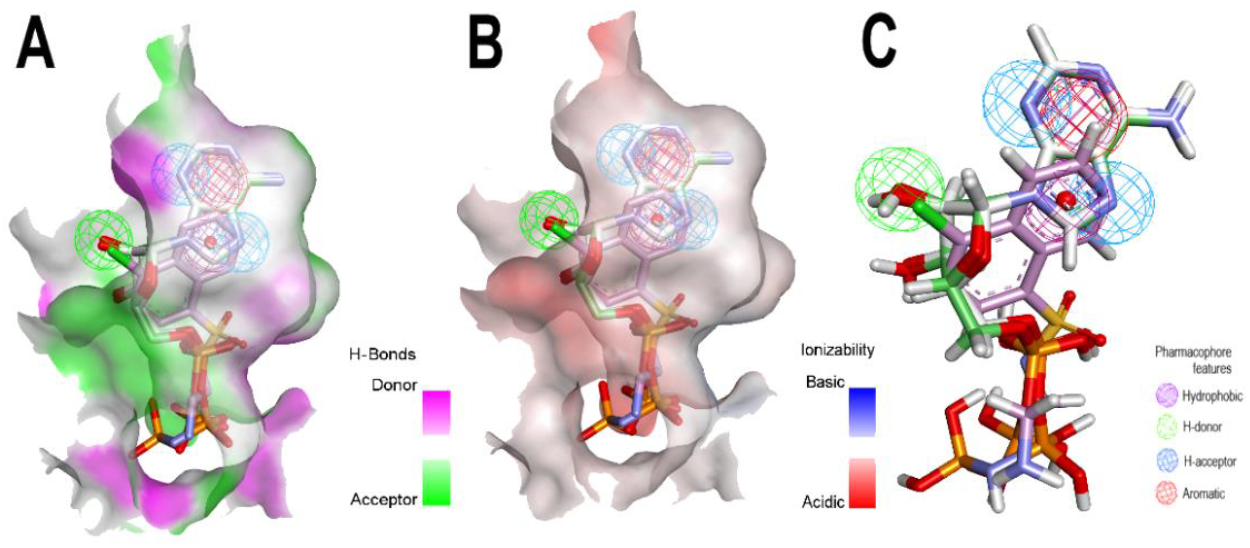
3D-representation of the NBS characteristics of EfAPH(3')-IIIa and pharmacophore model. A) H-bond surface and B) Ionizable surface of 3TM0. C) Target alignment of co-crystallized compounds highlighting the query features.

Subsequently, a high-throughput screening using the ZINC20 database containing 122,276,899 conformers were conducted. After a subsequent minimization and filtering of hits, considering mRMSD < 2.0, the results generating a total of 4,620 non-duplicate compounds that met the query features. Considering that molecular docking is highly sensitive to the conformation of the ligand binding site on the receptor ^43^, the next step after pharmacophore-assisted screening was to submit the 4,620 compounds to an ensemble docking protocol using PLANTS software.

To validation of virtual screening using the PLANTS were conducted redocking of each co-crystallized ligand in their respective receptor. For the first crystal, 3Q2J, the redocking parameters of CKI-7 showed an RMSD of 1.7 Å, Fig 2A. On the other hand, the redocking of ADP in the active site of 2BKK obtained an RMSD of 1.3 Å and for ANP in 3TM0 it was 0.6 Å as shown in Fig 2B and C, respectively. These redocking results are acceptable, especially considering the number of rotatable bonds presented in ADP and ANP co-crystalized molecules ^44^. The results of the virtual screening showed that the 4,620 compounds had binding affinities ranging from -140.1 to -42.1. In addition, the Kendall correlation comparing the database screening shows values of 0.1 between crystals 2BKK and 3Q2J, 0.11 between 2BKK and 3TM0 and 0.65 between 3Q2J and 3TM0, Fig. 2D. To select the best potential inhibitors of EfAPH(3')-IIIa the exponential consensus strategy ^45^ based on the docking scores with the three crystals was used.

**Fig 2.**
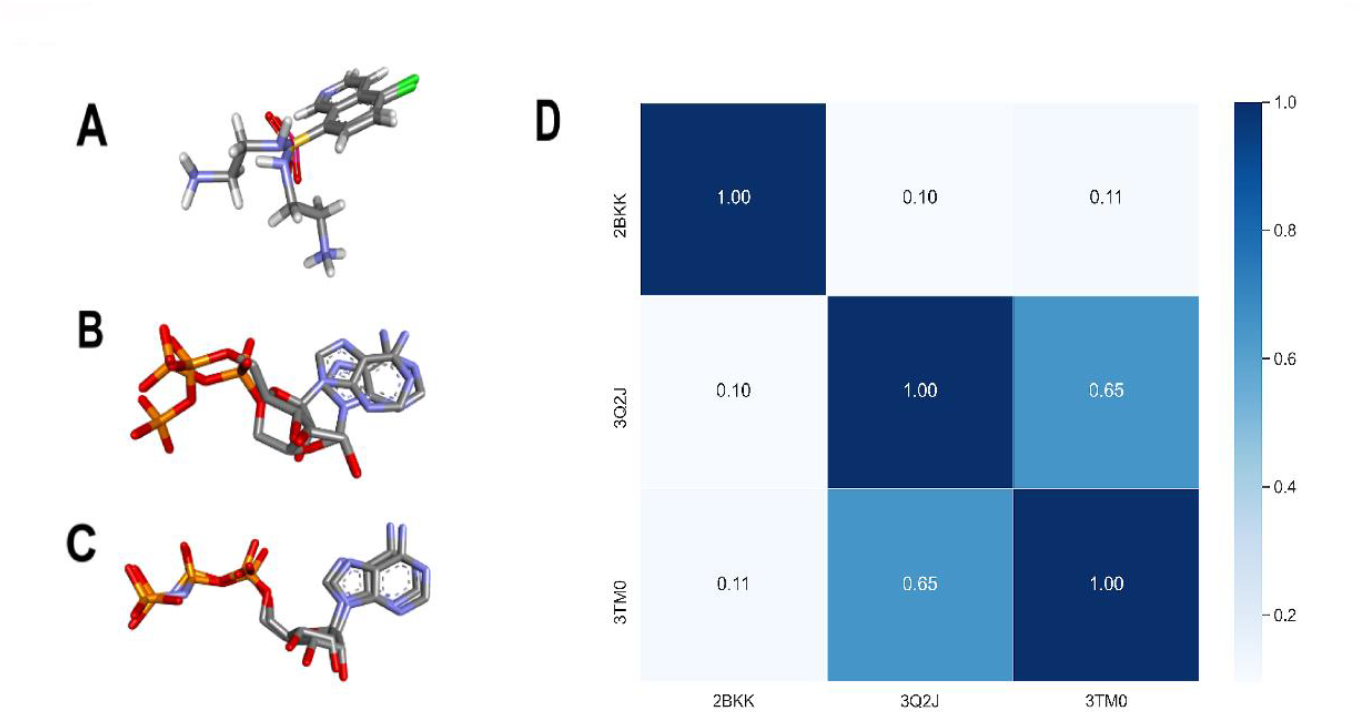
Redocking of co-crystals using PLANTS software and Kendall correlation of ensemble docking. A) Redocking of CKI-7 in the NBS of 3Q2J. B) Redocking of ADP in the NBS of 2BKK. C) Redocking of ANP in the NBS of 3TM0. D) Kendall correlation of 4,620 compounds docked in the three crystals.

The five molecules with the best ranking were selected for further analysis. In descending order of best-score (BS) were BS-1 (ZINC000104298353, (2Z,4E)-N-[2-[[(2S,3R,4S,5S,6R)-2-[(1R)-1,2-dihydroxyethyl]-4,5-dihydroxy-6-(7H-purin-6-ylamino)oxan-3-yl]amino]-2-oxoethyl]tetradeca-2,4-dienamide, Fig 3A), BS-2 (ZINC000828691112, 3-(4-{9-[3,3-bis(hydroxymethyl)cyclobutyl]-9H-purin-6-yl}piperazin-1-yl)-3-oxopropanenitrile, Fig 3B), BS-3 (ZINC000029546074, [(2R,3S,4R,5R)-5-(6-aminopurin-9-yl)-3,4-dihydroxyoxolan-2-yl]methyl N-[(2S)-2-amino-5-(diaminomethylideneamino)pentanoyl]sulfamate, Fig. 3C), BS-4 (ZINC000062624753, [[(2S,3S,4R,5S)-5-(3-acetylpyridin-1-ium-1-yl)-3,4-dihydroxyoxolan-2-yl]methoxy-hydroxyphosphoryl] [(2R,3R,4S,5R)-5-(6-aminopurin-9-yl)-3,4-dihydroxyoxolan-2-yl]methyl hydrogen phosphate, Fig. 3D), and BS-5 (ZINC000214484974 (1S,2S,3R,5S)-3-[7-[[(1R,2S)-2-(3,4-difluorophenyl)cyclopropyl]amino]-5-[(R)-propylsulfinyl]triazolo[4,5-d]pyrimidin-3-yl]-5-(2-hydroxyethoxy)cyclopentane-1,2-diol, Fig 3E).

**Fig 3.**
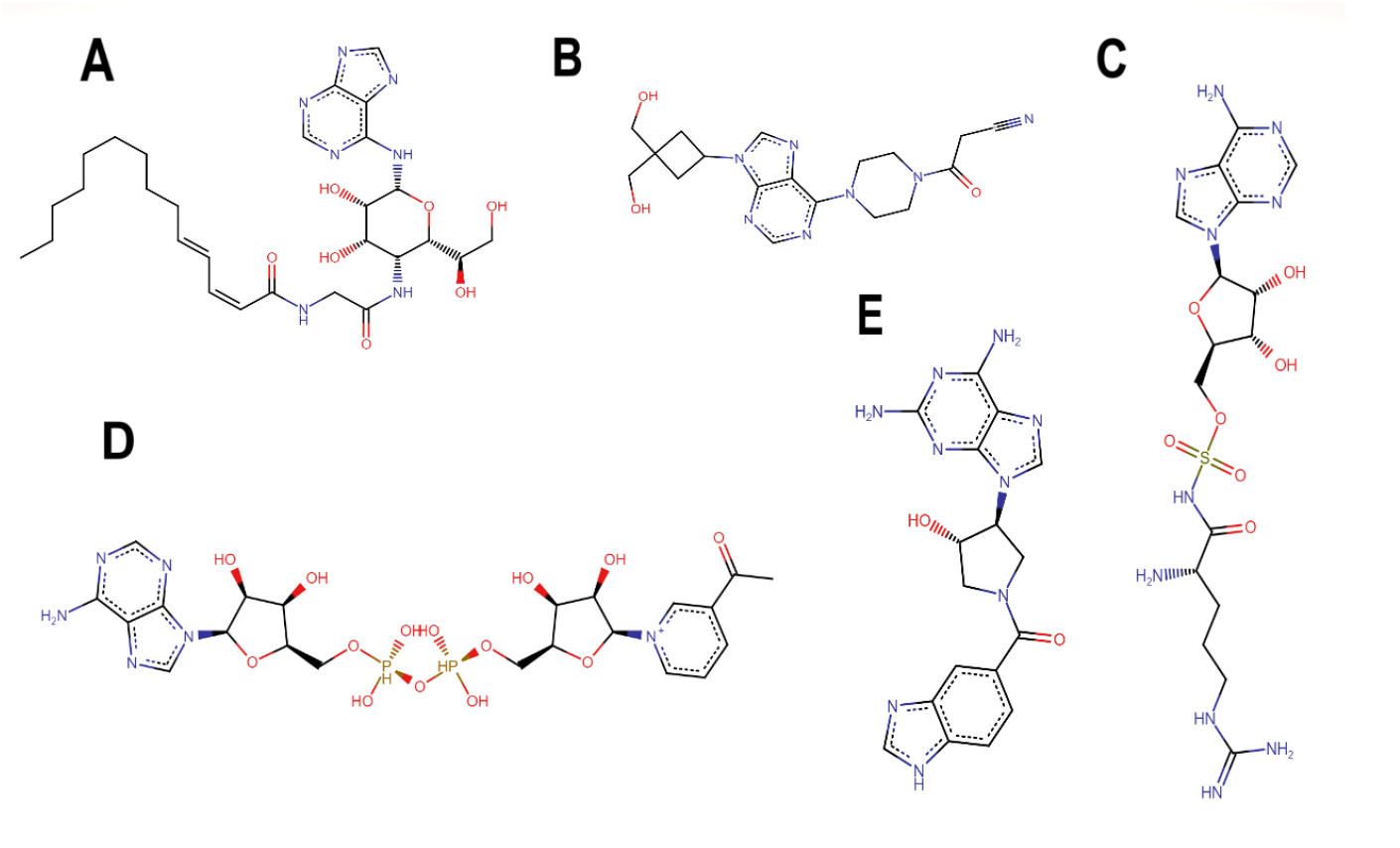
Structure of the best-scored nucleotide analogues on the nucleotide-binding site of EfAPH(3')-IIIa. A) BS-1 (ZINC000104298353). B) BS-2 (ZINC000828691112). C) BS-3 (ZINC000029546074). D) BS-4 (ZINC000062624753) and E) BS-5 (ZINC000214484974).

### 3.2 Pharmacokinetic and toxicological predictions of best-scored nucleotide analogues

*Analysis of the pharmacokinetic profile using the SwissADME server allowed us to observe that all nucleotide analogues except BS-2 have a low probability of being absorbed via the gastrointestinal tract and none of them are permeable to the blood-brain barrier. Additionally, BS-1 and BS-2 are potential P-gp substrates. However, only BS-4 has been shown to act as a potential cytochrome (CYP3A4) inhibitor*.

The results show that, in general, all the compounds have a promising profile due to their predicted inactivity in most of the parameters analyzed, Table 1. However, compounds BS-1 and BS-5 show immunotoxic and carcinogenic potential respectively, which compromises them as potential agents that could act as synergistic compounds of aminoglycoside activity. Based on these results, only BS2, BS3 and BS-4 will be further evaluated.

**Table 1.** Comparative pharmacokinetic profile of BS-2, BS-3 and BS-4. Pharmacokinetic analyses were performed with the SwissADME web server.

|  | Pharmacokinetic analyses |  |  |  |  |
| --- | --- | --- | --- | --- | --- |
|  | BS-1 | BS-2 | BS-3 | BS-4 | BS-5 |
| GI absorption | Low | Low | Low | Low | Low |
| BBB permeant | No | No | No | No | No |
| P-gp substrate | Yes | No | No | No | No |
| CYP1A2 inhibitor | No | No | No | No | No |
| CYP2C19 inhibitor | No | No | No | No | No |
| CYP2C9 inhibitor | No | No | No | No | No |
| CYP2D6 inhibitor | No | No | No | No | No |
| CYP3A4 inhibitor | No | No | No | Yes | No |

**Table 2.** Comparative toxicological profile of BS-2, BS-3 and BS-4. Toxicological analyses were performed with the Pro-Tox II web server. Values in parentheses refer to probability.

| Toxicological analyses |  |  |  |  |  |
| --- | --- | --- | --- | --- | --- |
| Properties | BS-1 | BS-2 | BS-3 | BS-4 | BS-5 |
| Hepatotoxicity | Inactive<br>(0.65) | Inactive<br>(0.60) | Inactive<br>(0.57) | Inactive<br>(0.65) | Inactive<br>(0.65) |
| Carcinogenicity | Inactive<br>(0.64) | Inactive<br>(0.64) | Inactive<br>(0.51) | Inactive<br>(0.66) | **Active**<br>**(0.50)** |
| Immunotoxicity | **Active**<br>**(0.99)** | Inactive<br>(0.91) | Inactive<br>(0.74) | Inactive<br>(0.99) | Inactive<br>(0.97) |
| Cardiotoxicity | Inactive<br>(0.71) | Inactive<br>(0.70) | Inactive<br>(0.79) | Inactive<br>(0.84) | Inactive<br>(0.81) |
| Mutagenicity | Inactive<br>(0.51) | Inactive<br>(0.77) | Inactive<br>(0.67) | Inactive<br>(0.64) | Inactive<br>(0.64) |
| Cytotoxicity | Inactive<br>(0.77) | Inactive<br>(0.64) | Inactive<br>(0.65) | Inactive<br>(0.85) | Inactive<br>(0.74) |

### 3.3 Interactions between the best-scored nucleotide analogues with EfAPH(3')-IIIa

To better understand the interaction of the three best-profiled compounds, the best pose complexed with the NBS of EfAPH(3')-IIIa was analyzed, and it was observed that there is a good correspondence of pose between these compounds and also the co-crystalized ANP, Fig 4A, similarly to what was previously reported ^40^.

**Fig 4.**
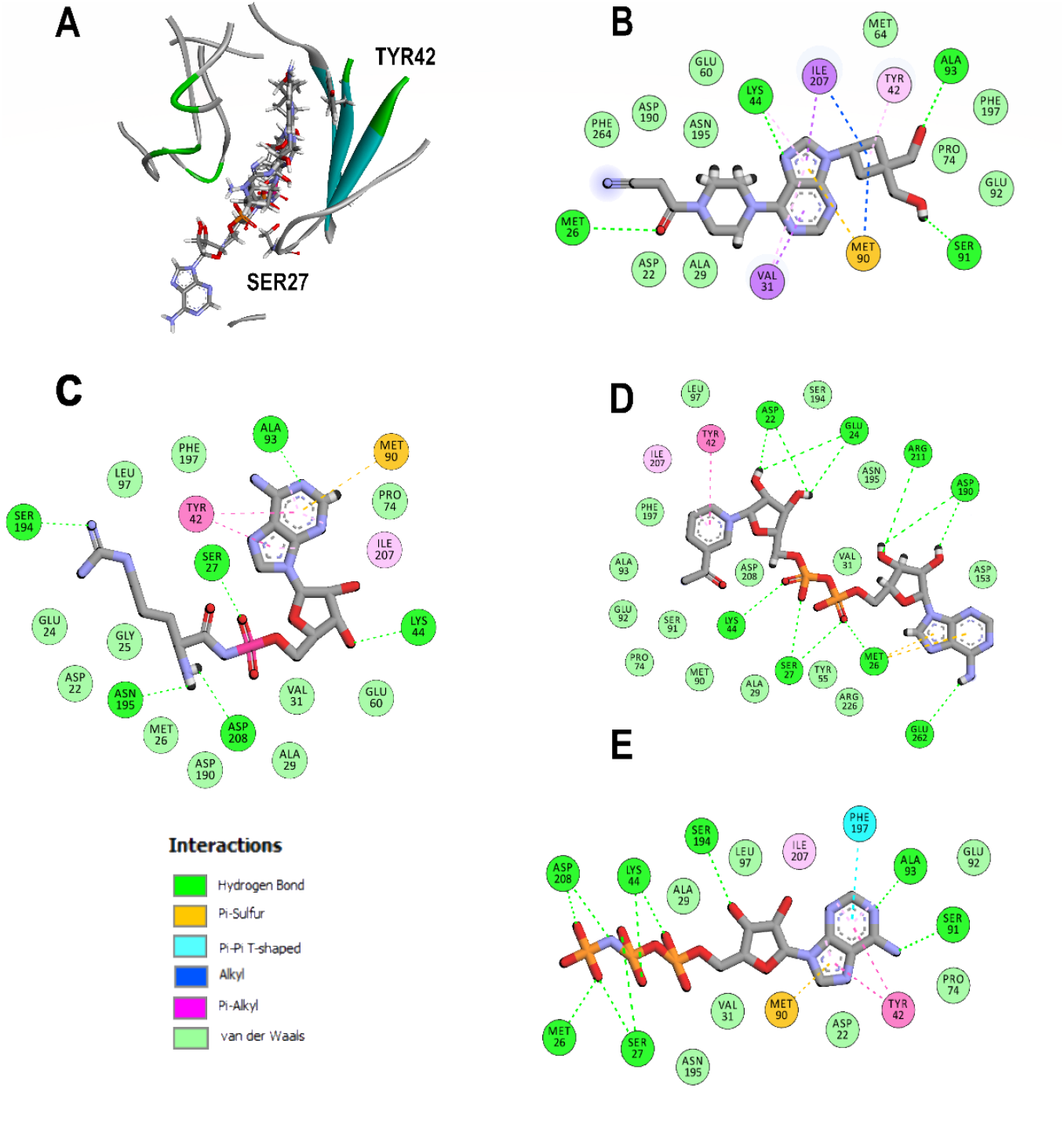
Analysis of the docked poses of the BS-2, BS-3, BS-4 and the co-crystalized ANP in the NBS of 3TM0. A) Sobreposition of the best-scored nucleotide analogues. B)2D interaction diagram of BS-2, C) BS-3, D) BS-4 and E) ANP.

Additionally, the representation also permits the observation that the NBS of 3TM0 complexed with the three ligands and the co-crystal reveal interactions with common residues like SER27 and TYR42. Notably, this last amino acid plays an essential role in the assistance of catalysis of the residue ASP190, and the formation of this same type of interaction with the adenine moiety of their physiological nucleotide ^28^.

Looking at the profile of the interactions in particular way, the cyclobutyl, adenine and pyridium rings of BS-2, BS-3, and BS-4 respectively face the aromatic ring of TYR42 resulting in the formation of pi-alkyl and pi-pi stacked interactions, Fig 4B-D. In addition, BS-1 also establishes hydrogen bonds between its ketone of the cyanoacetamide group with MET26, its imidazole group of adenine with LYS44 and hydroxy groups of the cyclobutyl ring with the residues of SER91 and ALA93, Fig 4B. At the same time, BS-2 establishes hydrogen bonds between SER27, LYS44, ALA93, SER194, ASN195 and ASP 208 respectively with their sulfur, 4'-hydroxyl of ribose, adenine, methylamine and formamidine groups Fig 4C.

On the other hand, BS-4 forms hydrogen bonds with the residues of ASP22, GLU24, ASP190, ARG211 through their 3/4'-hydroxyl of ribose, between SER27, MET26 and LYS44 of the alpha- and beta-phosphate groups and additionally between GLU262 and the amino group of adenine, Fig 4D. With regard to ANP, the interactions established were hydrogen bonds with MET26, SER27, SER91, ALA93, SER194, ASP195, ASP208 and additionally a pi-alkyl interaction with TYR42 (Fig. 4E).

## 4. Conclusion

In the approach adopted in this work, pharmacophore modeling was carried out followed by a first round of virtual screening against the ZINC20 database. A second virtual screening approach based on ensemble-docking to evaluate the affinity of the ligands resulting from the first screening was conducted using three different EfAPH(3')-IIIa crystals. The most promising ligands were ZINC000104298353, ZINC000828691112, ZINC000029546074, ZINC000062624753 and ZINC000214484974. Of these, the best toxicological and pharmacokinetic profiles were of ZINC000828691112, ZINC000029546074, ZINC000062624753. These potential inhibitors were able to interact with key amino acids such as MET26, SER27 and TYR42. Taken together, these results show that the nucleotide analogs studied in this work may represent a promising structure to interact with the nucleotide binding pocket of EfAPH(3')-IIIa. Our efforts will be focused on carrying out complementary analyses such as molecular dynamics, free energy calculations, among others.

